# Rapid, inexpensive measurement of synthetic bacterial community composition by Sanger sequencing

**DOI:** 10.1101/313932

**Authors:** Nathan Cermak, Manoshi Sen Datta, Arolyn Conwill

## Abstract

Simple synthetic bacterial communities are powerful tools for studying microbial ecology and evolution, as they enable rapid iteration between controlled laboratory experiments and theoretical modeling. However, their utility is hampered by the lack of fast, inexpensive, and accurate methods for quantifying bacterial community composition. For instance, while next-generation amplicon sequencing can be very accurate, high costs (>$30 per sample) and turnaround times (>1 month) limit the nature and pace of experiments. Here, we introduce a new approach for quantifying composition in synthetic bacterial communities based on Sanger sequencing. First, for a given community, we PCR-amplify a universal marker gene (here, the 16S rRNA gene), which yields a mixture of amplicons. Second, we sequence this amplicon mixture in a single Sanger sequencing reaction, which produces a “mixed” electropherogram with contributions from each community member. We also sequence each community member’s marker gene individually to generate “individual” electropherograms. Third, we fit the mixed electropherogram as a linear combination of time-warped individual electropherograms, thereby allowing us to estimate the fractional amplicon abundance of each strain within the community. Importantly, our approach accounts for retention-time variability in electrophoretic signals, which is crucial for accurate compositional estimates. Using synthetic communities of marine bacterial isolates, we show that this approach yields accurate and reproducible abundance estimates for two-, four-, and seven-strain bacterial communities. Furthermore, this approach can provide results within one day and costs ~$5 USD per sample. We envision this approach will enable new insights in microbial ecology by increasing the number of samples that can be analyzed and enabling faster iteration between experiments and theory. We have implemented our method in a free and open-source R package called CASEU (“Compositional Analysis by Sanger Electropherogram Unmixing”), available at https://bitbucket.org/DattaManoshi/caseu.

## Introduction

Model microbial communities – comprised of a small number of pre-defined, culturable taxa – are emerging as powerful tools in microbial ecology and biotechnology. Unlike wild microbial communities, whose underlying design principles are often obscured by complex environmental conditions and thousands of microbial “parts”, simple synthetic consortia can be studied precisely under controlled laboratory conditions. Through this approach, numerous studies have uncovered principles of microbial community interactions, assembly, organization, and evolution^1–8^. Furthermore, simple synthetic consortia hold great promise for biotechnology^9^, including synthesis of natural products that would be difficult to achieve with a single species^10^.

Despite the importance of model microbial communities, characterizing their composition (the proportional abundances of their constituent strains) quickly and cheaply remains challenging, since most standard methods have significant drawbacks (Table 1). On one hand, counting individual cells through colony formation on agar plates or with fluorescent labeling and flow cytometry is both cost- and time-effective and provides a direct measurement of population size. However, these methods can only be applied when strains are morphologically distinct or genetically tractable. On the other hand, next-generation sequencing can provide precise abundance estimates for arbitrary microbial communities, regardless of their composition, but it is typically very expensive and can take weeks to months to receive results.

**Table 1.** Comparison of methods for determining strain composition in simple model microbial communities. Sanger sequencing prices and turnaround times were obtained from Genewiz^14^.

| Method | Cost considerations | Speed | Measurement uncertainty | Biological limitations |
| --- | --- | --- | --- | --- |
| Next-generation sequencing (NGS) of marker gene amplicons | Entirely outsourced<br>>\$20/sample for library prep and sequencing<sup>11</sup><br>In-house<br>>\$1500/lane at university facility, plus library prep costs (\$37.5/sample)<sup>12</sup> | Typically weeks, sometimes months to get results<sup>13</sup> | Ideally limited by Poisson (counting) error. Given 50,000 reads, can detect members with abundance <0.01% | Requires marker gene that has unique sequence but conserved primer sites for all strains (e.g., 16S rRNA gene) |
| qPCR of marker genes | <\$1/sample for PCR | Same day | Large dynamic range but low accuracy | Similar to NGS. Requires designing specific primers or probes for each strain. |
| Plate counts of CFUs | Low (requires only agar plates) | Typically 2-3 days, depending on growth rates | Ideally limited by Poisson error | Strains must produce morphologically distinct colonies. Communities must be dissociable to single cells. |
| Fluorescent labeling of cells | Flow cytometer or microscope use | Same day | Ideally limited by Poisson error | Requires genetically tractable strains and spectrally distinct labels for each strain, potentially limiting communities to a few strains. |
| CASEU (this work) | \$4-6/sample for sequencing, plus <\$1/sample for PCR | As fast as next day | Fractional abundance error typically one percentage point | Similar to NGS for smaller consortia. |

Sanger sequencing has long been a cheap and effective method to characterize the taxonomy of bacterial strains in isolation, often by sequencing the 16S rRNA gene. This process typically begins by PCR-amplifying the 16S rRNA gene(s) from a pure bacterial culture containing a single strain. The result is a homogeneous pool of 16S rRNA amplicons (unless the strain has multiple copies of the 16S rRNA gene). Subsequently, the amplicon pool is subjected to a linear amplification process that yields DNA segments of different lengths^15^, where all segments of a given length have a fluorescent color label corresponding to the final (3’) base^16^. Then, DNA segments are sorted by length via capillary electrophoresis^17^, and the nucleotide sequence is determined from the corresponding sequence of fluorescent colors. Data are produced in the form of an electropherogram, in which fluorescent signal is plotted as a function of electrophoretic time (roughly corresponding to sequence position). Once characterized, the 16S rRNA gene sequence is often used as a taxonomic marker for a bacterial isolate.

In multi-strain bacterial communities where each member has a distinct 16S rRNA sequence, Sanger sequencing can be extended to characterize the presence and/or fractional abundance of each community member. The full complement of 16S rRNA genes present within a multi-strain community can also be PCR-amplified (typically with degenerate universal primers) and analyzed via Sanger sequencing. This process results in a “mixed” electropherogram. Like the single-strain electropherogram, a mixed electropherogram records the fluorescent signal as a function of electrophoretic time, but it now includes contributions from each of the strains present. Two approaches to characterize multi-strain community composition from mixed electropherograms have been developed previously (described below). However, unlike the new method we propose here, both prior approaches sought to characterize community composition without any prior knowledge of which strains were present.

In the first method^18^, a novel base-calling method was developed to preserve ambiguity at positions where multiple nucleotides were present, thereby allowing the authors to enumerate every possible constituent sequence. They then compared possible sequences to a database of known 16S rRNA gene sequences. Using this method, they reliably identified the bacteria present in numerous two- and three-species mixtures, including clinical samples^19–21^. However, this approach has not been used for quantification of strain abundance, and it is unclear how accurately the members of more complex communities (>3 strains) can be resolved.

In the second method^22^, the authors developed an algorithm to find a sparse set of strains whose combined DNA would be expected to generate the observed mixed electropherogram. To do this, they first created a database of predicted electropherograms (based on a statistical model of how gene sequences determine electropherograms) for 16S rRNA sequences of nearly 20,000 bacterial strains. They then used computationally solve for a small set of strains that could best reproduce the observed electropherogram. Applying this method to a mixture of five equally abundant strains, they detected at least eight strains, of which seven were closely related to strains in the actual mixture. However, their estimates of fractional abundances were noisy, varying from 5%-15% when the actual abundance was 20% each.

Here we develop and evaluate a new and distinct method for analyzing Sanger sequencing traces from amplicon mixtures as a fast (1 day) and inexpensive (~$5/sample) method for quantifying the fractional abundance of individual strains within simple model communities. It differs from previous approaches in two main ways. First, it assumes that one knows the full set of strains that might be in the mixture and experimentally measures their individual Sanger electropherograms. For model systems consisting of cultured isolates, this requirement is easily fulfilled. Second, our method accounts for a common mode of run-to-run variability not previously accounted for, which we show is necessary for accurate compositional estimates. We benchmark this method with multiple 2-, 4-, and 7-member communities of marine bacterial isolates, achieving a root-mean-square error of less than 1% and yielding results similar to next-generation sequencing. We also demonstrate the utility of this method by quantifying time dynamics of five model communities over two weeks. Overall, given its accuracy and broad applicability, we believe that this method will enable experiments with a wide range of simple synthetic microbial communities that were previously time- or cost-prohibitive.

We have also implemented our method in a free and open-source package for the open-source language R^23^, under the name “CASEU” for Community/Compositional Analysis via Sanger Electropherogram Unmixing. We provide functions for fitting and evaluating fit quality, both via the R language/terminal and through a graphical user interface.

## Approach

Our approach is to fit mixed-strain electropherograms as linear combinations of time-warped single-strain electropherograms. For a model bacterial community in which all component strains are known, it is possible to measure its mixed electropherogram, as well as each single-strain electropherogram. We thus sought to find a function relating the two that would allow us to extract the relative proportions of individual strains in the mixed electropherogram.

### A simple linear model is insufficient due to retention-time variability

Naively, it is reasonable to fit a mixed Sanger electropherogram as an abundance-weighted linear combination of single-strain electropherograms. However, this approach yields poor fits due to between-sample and within-sample variability in the run speed – that is, the rate at which molecules migrate during electrophoresis. This phenomenon, referred to as “retention-time variability”, is a well-known confounding factor in electrophoretic methods^24,25^ including Sanger sequencing. Indeed, we observed substantial retention-time variability in our measurements: technical replicates of the same sample sequenced on different days were often temporally offset from each other (by roughly ±1 base) and were sometimes stretched or contracted relative to one another by ±0.3% (see Figs. S1 and S2).

Instead, our fitting procedure conceptually involves two components: time-warping, which accounts for retention-time variability, and fitting a linear model. First, we warp (locally shift and stretch or contract) the time axis of single-strain electropherograms (Fig. 1). Second, we estimate strain abundances by fitting the mixed electropherogram as a linear combination of time-warped single-strain electropherograms. In practice, we do these steps simultaneously, by identifying warping parameters and abundance fractions that minimize the sum-of-squares difference between the observed and model-predicted mixed electropherogram, as follows:

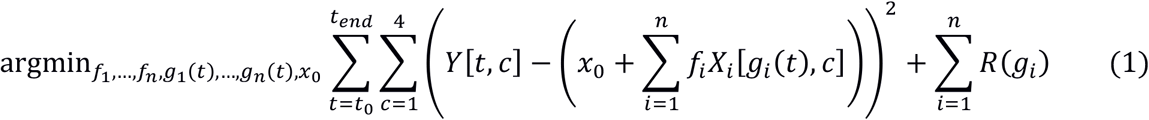

where

- *f_i_* is the abundance of strain *i*, where *i* ranges from 1 to *n*;
- *t* is an index of time, ranging from *t* _5_ to *t*_0_ *t_end_*;
- *Y* [*t*,*c*] is a matrix of the mixed electropherogram, with one row per timepoint and one column for each of the four fluorescence channels, *c*;
- *X_i_* [*t*,*c*] is a matrix of strain *i*’s electropherogram, with one row per timepoint and one column for each of the four fluorescence channels, *c*;
- *x*_0_ is a scalar accounting for constant background fluorescence;
- *g_i_* (*t*) is a warping function for strain *i*; and
- *R* (*g_i_*) is a quadratic penalty function for shifting and stretching individual electropherograms.

Because it is not possible to have negative abundances, we constrain strain abundances to be non-negative (*f_i_* > 0) by using non-negative least-squares fitting^26^.

**Fig. 1.**
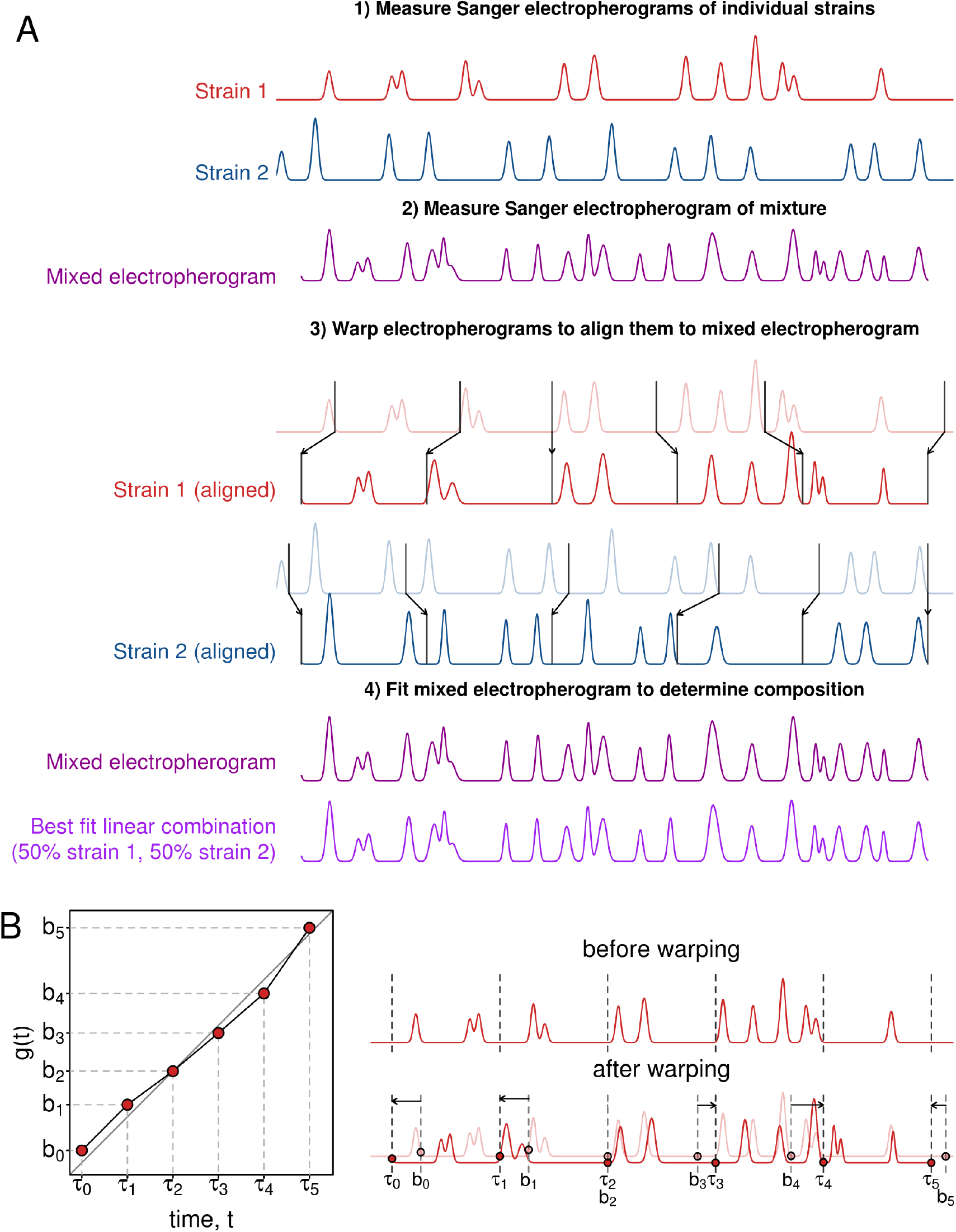
CASEU quantifies the fraction of individual strains in mixed communities by fitting mixed Sanger electropherograms as linear combinations of time-warped single-strain electropherograms. (A) Schematic of CASEU approach. Electropherograms shown are simulated for illustration purposes. For clarity, only a single fluorescence channel is illustrated. (B) Example of the continuous piecewise warping function used for alignment. The warping function is parameterized by six numbers (the values of the function at τ_0_, τ_1_, …, τ_5_). The figure shows an exaggerated warping with simulated electropherograms for illustration purposes.

### For time axis warping, we use a continuous piecewise time-warping function, which is sufficiently flexible to account for retention-time variability within an electropherogram

Within a given electropherogram, the relative run “speed” may vary substantially, such that certain sections are stretched, and others are contracted, compared to the average speed. To account for within-electropherogram variability, we use a continuous piecewise linear warping function *g_i_*(*t*) (see Fig. 1B), which divides the electropherogram into several segments, each of which can be locally stretched or contracted^24^. To prevent unreasonably large stretching or shifting, we include a small quadratic penalty for moving the end of each segment from its original location (*R*(*g_i_*) ∝ Σ*_j_*(*τ_j_* − *b*_j_)^2^ Where *τ_j_* and *b_j_* are as shown in Fig. 1B). To determine the optimal number of segments, we systematically varied the number of segments and aligned technical replicates to each other. We found that using five segments enabled us to align all samples precisely to either of their two technical replicates over a region of ~630 bases (Fig. S2). Using only a single segment yielded poor alignments between technical replicates (Fig. S2) and produced mediocre estimates of known mixture fractions (Fig. S3). Using more than five segments did not improve the alignments between technical replicates (Fig. S2), but increased computation time.

## Results

### Benchmarking algorithmic performance

#### To benchmark CASEU’s performance, we analyzed a series of mock bacterial communities of 2, 4, or 7 bacterial strains with known fractional abundances

We prepared these communities by PCR-amplifying the 16S rRNA gene from each single strain (here called “A” through “H”), and mixing together amplicons from different strains in known fractions. By analyzing mixtures of amplicons, rather than mixtures of cells, we could measure sequencing and algorithmic performance independent of biases due to DNA extraction efficiency, PCR efficiency, or 16S rRNA gene copy number. Using mock communities, we assessed the following metrics of algorithmic performance:

- accuracy of fractional abundance estimates, by systematically varying the abundance of community members between 1.3%-95%;
- reproducibility, by sequencing each sample three times on separate days;
- ability to differentiate closely related strains, by varying phylogenetic distance between strains; and
- ability to correctly reject the presence of “decoy strains”, which are included as potential community members in the fits but were absent in reality.

#### CASEU infers strain fractional abundances accurately and reproducibly in many 2-, 4-, and 7-strain communities, while rejecting the presence of decoy strains

We first analyzed two-strain mixtures for which the proportion of a single strain varied from 5% to 95% (Fig. 2). Across all two-strain communities (except the mixtures of strains A and B, see below), fractional abundance estimates were accurate with an average absolute deviation between the expected fraction and the observed fraction of 0.6 percentage points (range 0.05% to 1.8%). Furthermore, abundance estimates were consistent across independently sequenced technical replicates; the average standard deviation of triplicate measurements was 0.55 percentage points (range 0.06% to 0.93%).

**Fig 2.**
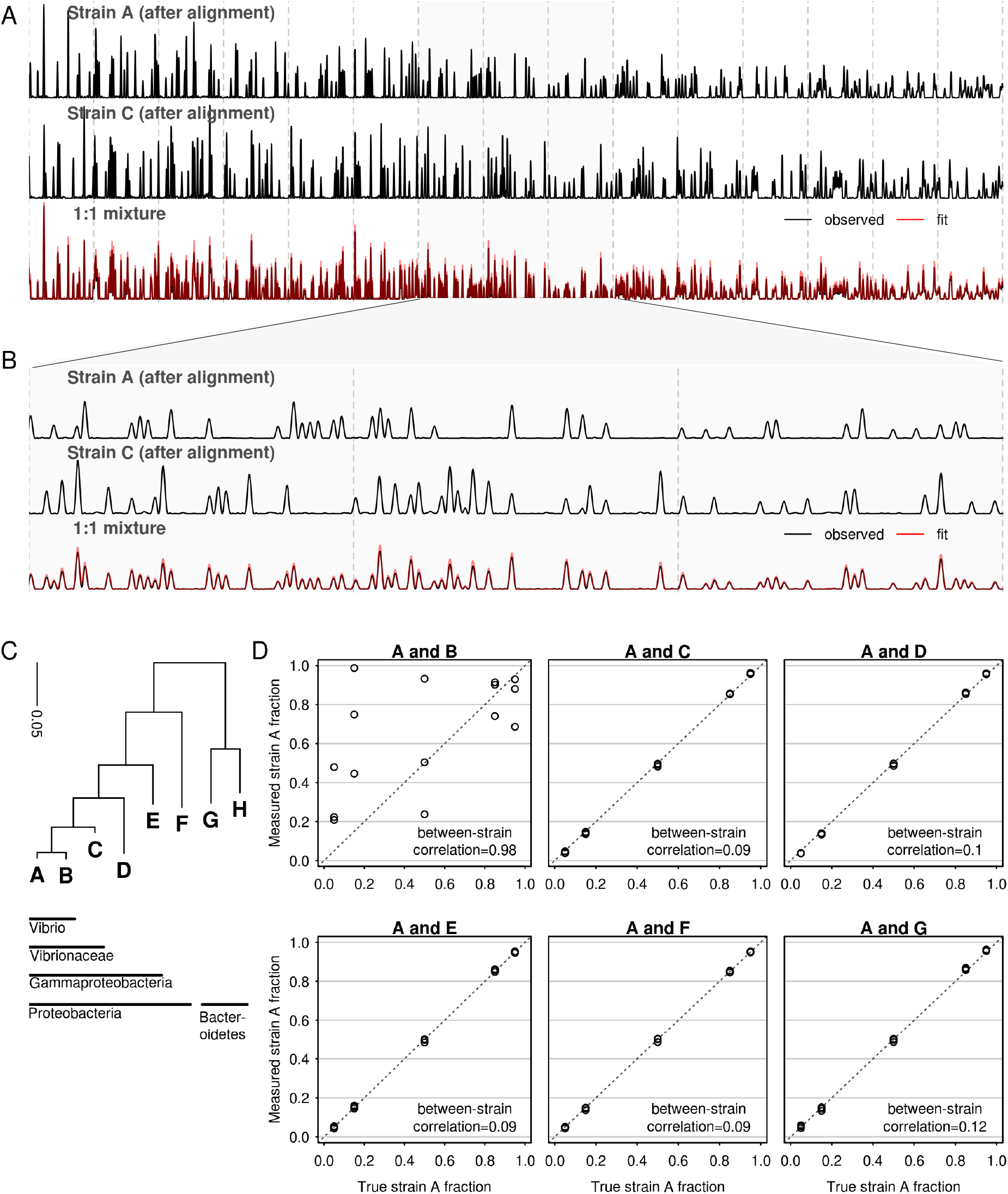
CASEU accurately resolves composition of two-strain mock communities. (A) An example alignment and fit over approximately 630 bases, showing a single fluorescence channel. Top and middle traces show reference electropherograms of individual strains (after warping). Bottom trace shows the electropherogram of a 1:1 mixture (black) and best-fit weighted sum of aligned references (red). (B) Zoom-in of segment of (A), showing alignments and fit over approximately 120 bases. (C) Phylogenetic tree of genes chosen for analysis, made using nearly full-length 16S sequences from ref 27. We aligned these sequences using the SINA Alignment Server^28^ and made an approximate maximum-likelihood tree using FastTree 2.1.10 with the default options^29^. (D) Estimated mixture fractions plotted against the true ratio at which the sequences were mixed (circles). These mixture fractions have been corrected for errors in stock concentration (uncorrected fraction data shown in Fig. S3, bottom row).

In larger communities (4- and 7-strain mixtures), fractional abundance estimates were similarly accurate, even for low-abundance community members. To test the effect of strain evenness on fractional abundance estimates, we prepared 4- or 7-strain communities whose strain abundances were distributed according to a power law (*f_i_* ∝ *i*^−*α*^), where we varied the value of the exponent *α*. This allowed us to assemble communities of varying evenness (Fig. 3), ranging from those in which all strains were at equal abundance (*α* = 0) to those in which the dominant strain was 50-fold more abundant than the least abundant strain (*α* = 2). Across these communities, abundance estimates were similarly accurate compared to the two-strain communities, with root-mean-square (RMS) errors of 0.73 and 1.14 percentage points (maximum errors of 1.9 and 3.9 percentage points), respectively for the 4- and 7-strain communities. Furthermore, the magnitude of error in a strain’s abundance was nearly independent of that strain’s abundance in the community (Fig. S5A). The standard deviations we observe between triplicate results were comparable to what would be attained by counting based methods (e.g. NGS or plate counts) with ~5200 counts (reads or colonies) per sample (Fig. S5B).

**Fig 3.**
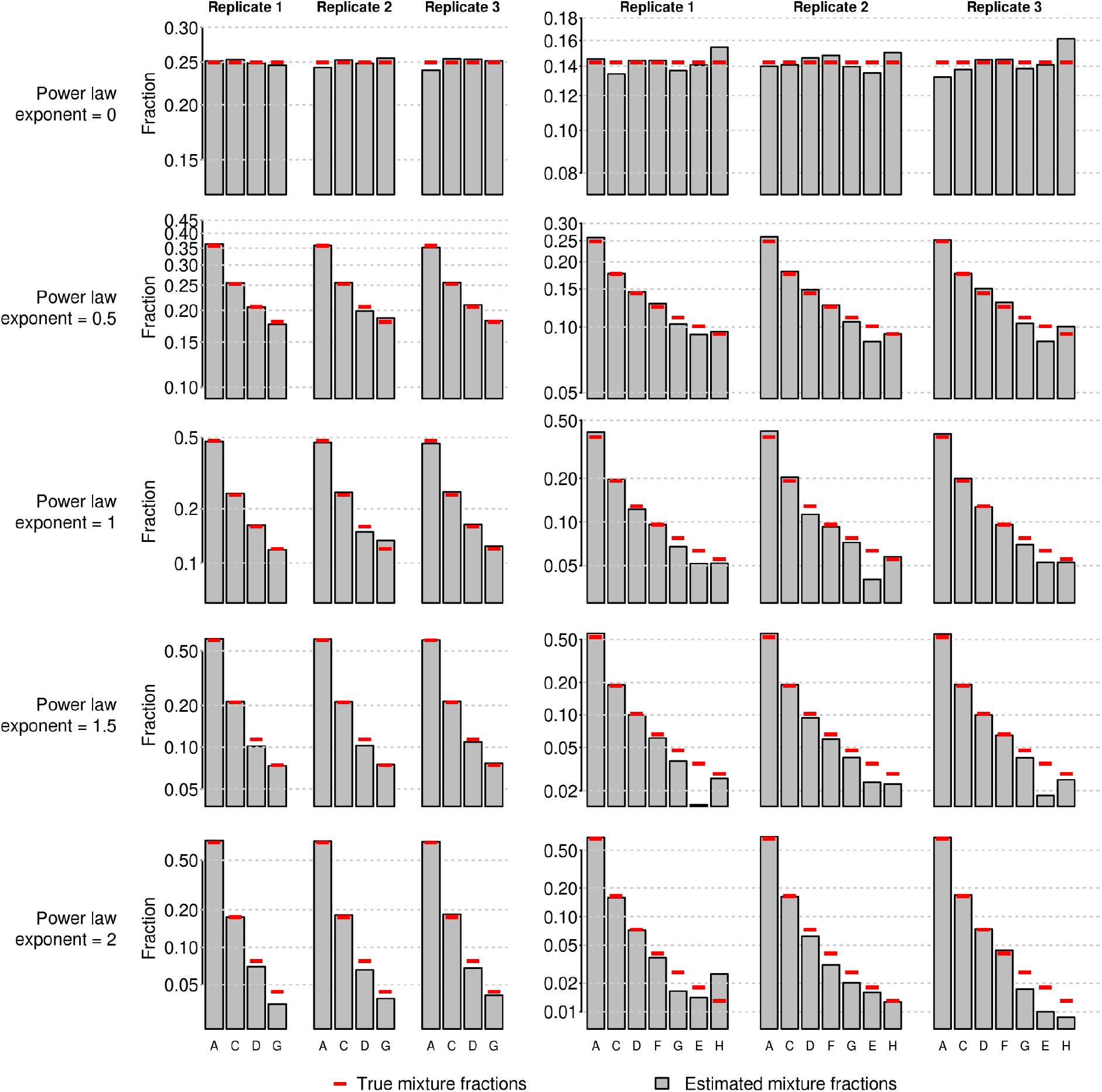
CASEU provides reliable estimates of community composition in mixtures of 16S amplicons from four (left) or seven (right) strains. Solid bars show measurements after accounting for stock concentration error (uncorrected data is given in Fig. S6), red lines show true mixture proportions based on power law distributions. In power law distributions, the abundance of the *i*^th^ most abundant strain is proportional to 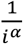 where *α* is the power law exponent.

It is important to not only estimate the abundance of a strain known to be present, but also to correctly determine when a strain is absent. To test whether CASEU is susceptible to erroneously finding strains that were not present, we re-fit all our two- and four-species communities, this time including all strains (except for B) as possible “decoy” community members. In nearly all cases, CASEU correctly rejected the presence of strains that were not included in the community (Fig. 4). Notably, CASEU erroneously found non-zero amounts of strain D in some samples where only strains A and C were present. We attribute this to the similarity between electropherograms of strains C and D (discussed below).

**Fig. 4.**
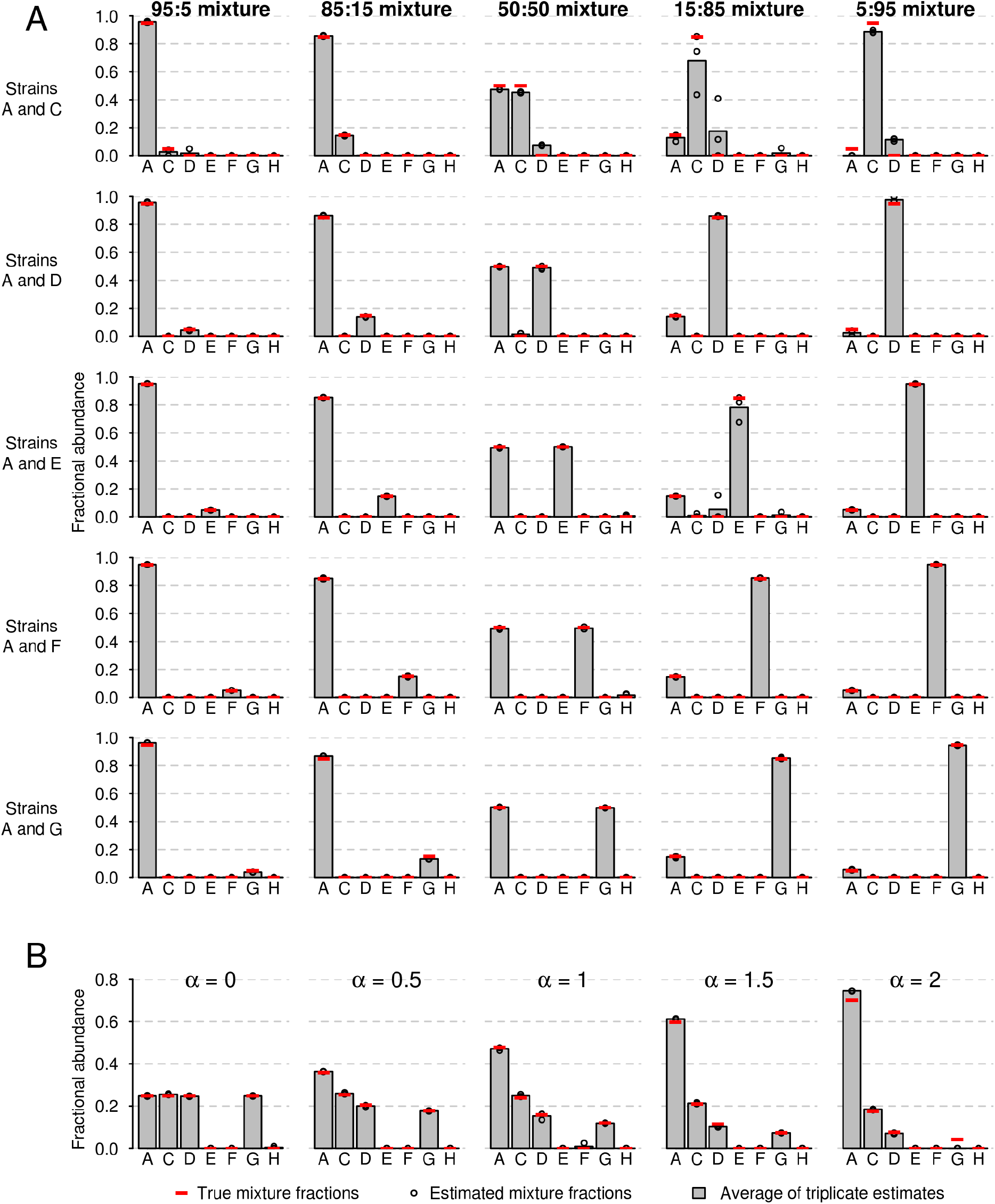
CASEU correctly infers the absence of strains that were not present in the mixture. (A) Community composition estimates of two-strain mixtures (as in Fig. 2D), in which five extra “decoy” strains were included as potential community members to test CASEU’s ability to infer strain absence. Bars indicate CASEU estimates (average of three replicates); open circles indicate each of the three replicate estimates, and red lines indicate the true values. Estimates are corrected for errors in stock concentrations. (B) Community composition estimates of four-strain mixtures (as in Fig. 3; *α* is the power law exponent), in which three extra “decoy” strains were included as potential members. Bars and points are as in (A).

#### To differentiate strains, CASEU requires that their electropherograms are dissimilar

We quantified similarity as the correlation between two electropherograms after aligning one to the other. Our mock communities contained mixtures of strains with varying degrees of electropherogram similarity, ranging from 0.98 (for strains A and B) to 0.09 (for strains A and G) over a 630-basepair region of the 16S rRNA gene (Fig. S4). While CASEU failed to differentiate strains A and B which have an average post-alignment correlation of 0.98 (Fig. 2), it accurately estimated fractional abundances for all other communities (Figs. 2 and 3) containing between-strain correlations of up to 0.82 (strains C and D; Fig. 3). However, strain D was sometimes mistakenly found in the mixtures of strain A and C, suggesting it may sometimes be mistaken for strain C. This suggests that strains with correlations of 0.82 or greater are unlikely to be clearly resolvable with CASEU. In our dataset, this corresponds to roughly within-genus distances or closer, but the relationship between CASEU resolvability and phylogeny may depend on the specific strains of interest.

### Evaluation on synthetic model communities

#### We envision CASEU as a rapid, cheap method for characterizing the structure and dynamics of simple synthetic microbial communities

To demonstrate this use case, we performed two experiments with seven four-strain model communities of unknown fractional composition.

1. **Paired compositional analysis of model communities**. We extracted DNA from each community, then amplified and sequenced each sample twice, once via Sanger sequencing (16S rRNA V1-V9 hypervariable regions) and once via next-generation Illumina sequencing (16S rRNA V4-V5 hypervariable regions) (Fig. 5). Importantly, this analysis does not compare the accuracy of the two methods, since the true fractional abundances are unknown, but rather whether their fractional abundance estimates are consistent.
2. **Time dynamics of serially transferred communities.** We maintained each model community in serial batch culture, passaged daily with a 100-fold dilution, and sampled them five times over two weeks for Sanger sequencing. We also used Illumina sequencing of the 16S V4-V5 region at the final time point to compare the consistency of fractional abundance estimates for the two methods.

**Fig. 5.**
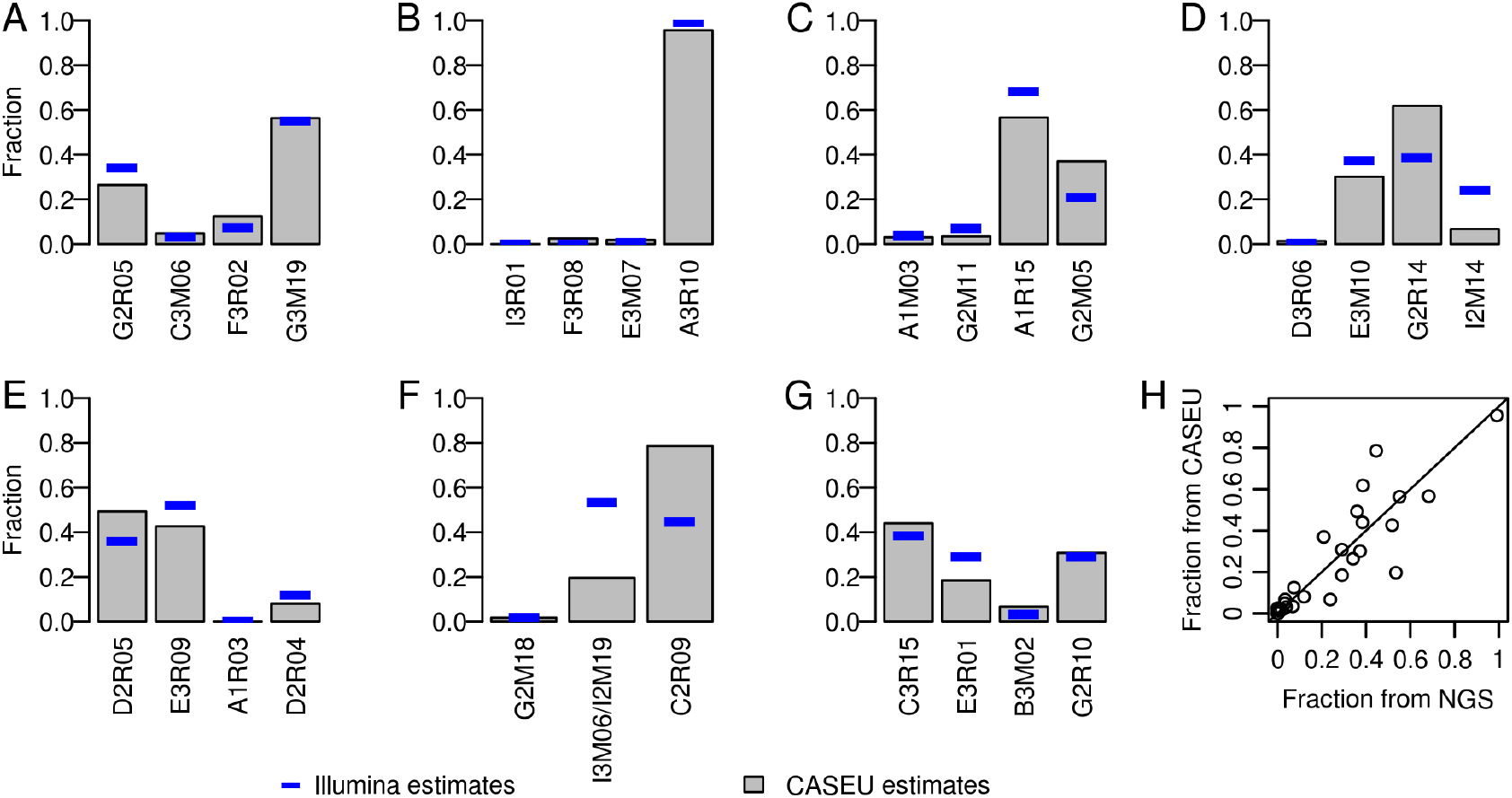
CASEU yields estimates of community composition that are typically consistent with Illumina 16S sequencing. (A-G) Bar plots indicate results of CASEU analyses of mixtures of saturated cultures of four bacterial strains. Solid blue lines show estimates obtained from Illumina sequencing of the 16S rRNA V4-V5 hypervariable region. (H) Fractional abundance estimates for CASEU vs Illumina sequencing. Solid line shows equality.

We also demonstrate two cases in which CASEU fails to infer fractional abundances correctly and produced a visibly poor fit. Assessing the quality of the fit allowed us to detect errors, which we could not have done through next-generation sequencing.

#### CASEU provides community composition estimates consistent with next-generation Illumina sequencing of 16S rRNA amplicons (Fig. 5), despite differences in library preparation procedure and sequencing technology

In particular, across all communities, fractional abundance estimates between the two methods were highly correlated (Pearson correlation 0.88, Fig. 5H). Furthermore, in five out of seven communities, we observed strong quantitative agreement between Illumina estimates and CASEU estimates, with an RMS difference of 6.9 percentage points (Fig. 5A,B,C,E,G). We observed a similar level of agreement for the final timepoints of serially transferred communities (RMS difference of 2.1 percentage points.)

#### Major discrepancies between CASEU and Illumina sequencing can be attributed to a single group of closely related strains

For two model communities, we observed sizable differences between the CASEU and Sanger estimates (Fig. 5D,F; RMS differences of 15 and 28 percentage points, respectively). These differences were driven by the abundances of three strains (I3M06, I2M14, and I2M19), whose electropherograms are too similar to be distinguished by CASEU. In both model communities, these strains were estimated by CASEU to be at substantially lower fractions than was estimated by next-generation sequencing. While we remain uncertain as to why these strains are detected less with CASEU than Illumina, it may be a result of CASEU and Illumina relying on different primers and amplification protocols.

#### CASEU can be used to quantify time dynamics of model communities

Using CASEU, we also quantified changes in community composition over time for seven model communities. We found that four communities rapidly reached an equilibrium state dominated by a single strain, with one strain exceeding 90% abundance within five days (Fig. 6B,C,D,E). However, one community continued to change in composition throughout the course of the experiment, suggesting that the community has not reached a stable equilibrium composition. Therefore, CASEU can identify coexistence and equilibrium states in model microbial communities.

**Fig 6.**
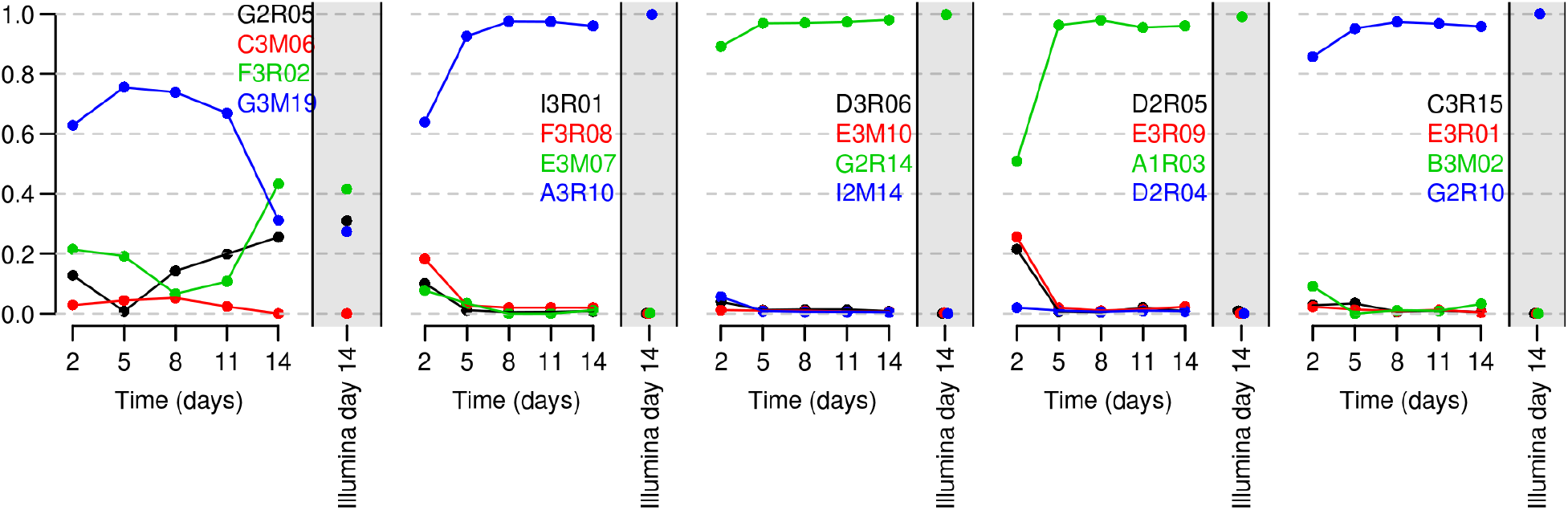
CASEU enables quantification of temporal dynamics in model four-strain communities. Temporal dynamics of five four-member communities measured over two weeks via CASEU, as compared to the final time point measured by Illumina sequencing of 16S rRNA V4-V5 amplicons.

#### Assessment of CASEU fits allows us to detect errors in sample preparation or sequencing

Across CASEU analyses performed with these four-strain model communities (39 in total), we identified two cases in which CASEU produced poor fits as quantified by the correlation between the observed and predicted traces. In the first case, the predicted electropherogram had a correlation of only 0.63 to the observed electropherogram, compared to >0.98 for other samples from the same model community. This poor fit alerted us to a low-quality Sanger sequencing electropherogram for one strain in the community, which contained a large anomalous fluorescence spike (Fig. S7B). In the second case, CASEU yielded a correlation between predicted and observed traces of 0.45, compared to >0.95 for other samples from the same model community. This poor fit was caused by the presence of a contaminating strain, which was not included in the fit (Fig. S7C). Including the contaminating strain increased the fit correlation to 0.95. Thus, while we only rarely observed poor fits, CASEU includes a simple metric that enables users to identify and exclude problematic samples.

## Discussion

In microbial ecology, model communities have emerged as a useful intermediate between single-species microbiology and complex natural communities. Here, we demonstrate that Sanger sequencing can be used for **rapid, inexpensive, and accurate** quantification of model community composition.

- **CASEU can provide rapid results.** Sanger sequencing requires a simple sample preparation protocol with a single PCR step, followed by outsourced Sanger sequencing. Therefore, the time to acquire results is largely limited by sequencing time, which is often less than one day. In contrast, next-generation sequencing requires a more time-consuming library preparation protocol, often with multiple PCR steps for adaptor ligation and barcoding. Furthermore, runtime for an Illumina MiSeq routinely exceeds one day (e.g., 40 hours for paired-end 150×150 sequencing), but can require weeks to months, if outsourced.
- **CASEU can be inexpensive.** Sanger sequencing has a fixed cost per sample (here, $4 USD for sequencing and roughly $1 for PCR and cleanup), whereas NGS has large base cost (typically more than $1,000 USD per sequencing lane), plus often tens of dollars per sample for library preparation, particularly if library preparation is outsourced.
- **CASEU is accurate for simple model communities.** Sanger sequencing provides an accurate and reproducible means to quantify composition for model communities, achieving similar results as NGS for model communities, and errors of less than one percentage point for mock communities.

To determine if CASEU is appropriate for your application, we recommend considering the following factors as they pertain to your model community.

1. **Number of strains.** Here, we demonstrate that CASEU can provide accurate fractional abundance estimates for communities of 2, 4, and 7 strains. However, CASEU may be suitable for larger model communities, as we did not identify an upper bound on the number of resolvable members.
2. **Resolvability of strains.** We found that strain resolvability depends on the correlation of their electropherograms, which is distinct from their aligned sequence similarity. Therefore, for your particular community, we recommend Sanger sequencing each individual strain and verifying that their electropherograms cannot be aligned to be highly correlated, which can be done with our R package.
3. **Low-abundance strains.** Given typical errors of 1-2%, CASEU cannot resolve community members at fractional abundances below 1%. If this dynamic range is needed, alternatives like qPCR or NGS may be more suitable.
4. **Sources of bias.** CASEU shares the same limitations of all DNA-based approaches for quantifying community composition, including bias in DNA extraction and amplification efficiency. Importantly, NGS, qPCR, and CASEU do not yield cell counts, but instead yield sequence abundance. While sequence abundance is expected to be roughly proportional to cell count for any given strain, this relationship may vary between strains depending on gene copy number, growth phase, DNA extraction efficiency, and amplification efficiency.

More broadly, we believe that our Sanger sequencing deconvolution approach can be extended beyond the 16S gene. For example, CASEU might be used with model communities containing closely-related strains by using other marker genes (e.g. Vibrio communities that are poorly resolved by 16S but easily differentiated by hsp60 sequences^30^), or even communities containing both fungal and bacterial members (for example, cheese rind model communities^7^) by amplifying both 16S and 18S or ITS sequences simultaneously through multiplexed PCR. Beyond microbes, CASEU might extended to quantify aneuploidy using marker sequences with conserved primer sites present on all chromosomes^31^. Overall, we believe CASEU provides a versatile tool to assess sequence-variant composition in multiple contexts.

## Acknowledgments

Thanks to Andrea Velenich and Jeff Gore for helpful discussions, and to Otto Cordero for generously sharing strains. AC acknowledges support from the Simons Collaboration on Principles of Microbial Ecosystems (PriME).

## Contributions

MSD and NC designed the model and fitting algorithm, performed the experiments with amplicon mixtures, and analyzed all Sanger sequencing data. AC performed experiments with synthetic model bacterial communities. AC and MSD analyzed Illumina sequencing data. MSD and NC wrote the paper with input from AC.

## Methods

### Preparing mixtures of 16S amplicons and sequencing

For two-, four-, and seven-strain mixtures, genomic DNA was extracted as previously reported^15^. 16S genes were amplified with 27F (AGAGTTTGATCMTGGCTCAG) and 1492R (TACGGYTACCTTGTTACGACTT) universal primers, as follows:

| Reagent | Volume |
| --- | --- |
| ddH<sub>2</sub>O | 23.5 µL |
| 5X HF Buffer | 10 µL |
| dNTPs (10mM) | 1 µL |
| 27F primer (3uM) | 5 µL |
| 1492R primer (3uM) | 5 µL |
| Phusion polymerase | 0.5 µL |
| Genomic DNA | 5 µL |
| Total | 50 µL |

PCR cycle conditions were as follows:

| Step | Temperature | Duration |
| --- | --- | --- |
| Initial denaturation | 98°C | 30 seconds |
| Amplification (30 cycles) | 98°C | 30 seconds |
|  | 50°C | 30 seconds |
|  | 72°C | 90 seconds |
| Final extension | 72°C | 10 minutes |

For experiments shown in Figures 2 and 3, we ran six PCRs for each strain to ensure that we had sufficient amplicon DNA to prepare all the mixtures. We pooled each set of six reactions into a single tube, then SPRI-cleaned the products. We estimated DNA concentrations via Nanodrop, and subsequently diluted all samples to 3 ng/μL (concentration measurements required subsequent computational correction, see Materials and Methods). We added 5 μL of 27F primer at 15 μM to 40 μL of amplicon at 3 ng/μL, to yield 45 μL with a primer concentration of 1.6 μM and a template concentration of 2.6 ng/μL. These concentrations are what Genewiz recommends for Sanger sequencing^32^. We split the 45 μL of sample into three separate plates, each with 15 μL of sample per well, and submitted each plate on a different day over the course of one week.

Sequencing was performed by Genewiz as a drop-off service for $6/sample (<48 samples) or $4/sample (>48 samples). We routinely received results within 24 hours of submitting our samples. Our ABIF file metadata suggests Genewiz sequencing was performed on a 3730xl DNA Analyzer, using BigDyeV3.

### Processing ABIF files

We used the ‘sangerseqR’ Bioconductor package^33^ in R^23^ to read in ABIF (.ab1) files. In ABIF files, there are two types of data we considered using: “raw” fluorescence traces, and “processed” data. While the details of the processing method are not available, the process appears to involve baseline subtraction, low-pass filtering, and an unknown temporal adjustment. Attempts to use the “raw” traces were stymied by the poor temporal alignment of the traces and required searching a much larger range of alignment parameters, yielding a significantly slower analysis. Before analysis, we additionally normalized the amplitudes of all reference files such that the mean amplitude was one over the region to be used for alignment.

### Algorithm for fitting mixed electropherograms

We initially tried optimizing Equation (1) via the Nelder-Mead algorithm (also called downhill simplex) but found that this method tended to yield solutions that were very dependent on starting estimates (suggesting many local minima). Instead, we adopted an approach in which we determine the warping parameters for one strain at a time. To determine the warping parameters for a single strain (“aligning a single strain”), we use the dynamic programming approach pioneered in correlation-optimized warping^24^. In brief, we first calculate the sum of squared errors over a 2D grid of values for parameters *b_1_* and *b_2_* (the boundaries of the first warping segment). (Note that to do so, we rapidly calculate optimal f_i_ values for each *(b_1_*, *b_2_*) pair using non-negative least squares.) For each possible value of *b_2_*, we only keep track of the best value of *b_1_* and its corresponding error. We then repeat that same process, but this time evaluating the error for a grid of possible values for *b_2_* and *b_3_* (the boundaries of the second segment). For every *b_2_*-*b_3_* pairing, we calculate the error over that segment, plus the lowest possible error for that value of *b_2_* over any value of *b_1_*. We then record the lowest error obtainable for any given value of *b_3_*, and the corresponding *b_2_* that yields that optimum. We repeat this process for all five warping segments. This process does not necessarily yield a globally optimal solution because, in order for the errors for each segment to be additive, we allowed each segment to have its own amplitude parameter *f_i_*. However, ultimately this parameter *f_i_*, must be the same for all segments. We thus globally refine parameter estimates by minimizing equation (1) directly via Nelder-Mead, starting from the (*b_1_-b_6_*) estimates obtained as described above (which are usually very near the final optimum).

To align multiple strains, we sequentially align strains one after another. We first align each strain individually, identify the one that yields the greatest improvement in the fit, and fix that strain’s alignment parameters (*b_1_*-*b_6_*). We then repeat that process for the remaining strains, always greedily fixing the parameters of whichever strain yields the greatest improvement in the fit (reduction in squared error). Our rationale was that this would ensure that we always fit the majority component of the mixture before fitting the minority components.

More formally, our algorithm is as follows:

**Inputs**:

- *Y*[*t,c*]-mixed electropherogram matrix
- *X*_1_[*t,c*], *X*_2_[*t,c*], …, *X_n_*[*t,c*] - Individual reference electropherogram matrices (*n* is number of strains)

**Run:**

1. Initialize *F* = Ø, the indices of strains for which the alignment parameters have been fixed.
2. Initialize *U* ≠ {1, …, *n*}, the indices of strains for which the alignment parameters have not yet been fixed.
3. Initialize *A* as an empty *n*×6 matrix for the alignment parameters.
4. While *U* ≠ Ø
  1. For every strain index *i* in *U* find a warping of *X_i_* via dynamic programming that yields the greatest reduction in equation (1), conditional on all strains with fixed alignments *X_f_* for *f* ∈ *F*
  2. Identify which strain *j*, yields the most improvement to the fit.
  3. Fine-tune the alignment parameters of strain *j* by downhill simplex.
  4. Record strain *j*’s alignment parameters in row *j* of matrix *A*.
  5. Add strain *j* to *F*, remove it from *U*
5. Fine-tune all alignment parameters simultaneously by optimizing over *A* via downhill simplex, starting from *A* (the best individual alignments obtained in step 4).

On a HP laptop with an Intel Core i7-7500U CPU and 16 GB RAM, our fitting approach took on average 10.8 seconds for a two-strain mixture, and 138 seconds to fit a seven-strain mixture. Notably, the computation time is expected to be at worst quadratic in the number of strains that could potentially be in the mixture.

### Accounting for concentration errors

Before mixing amplicons to make mock communities, we attempted to normalize all amplicon stock solutions to the same concentration, as measured by Nanodrop. However, the Nanodrop has limited precision, and as such we have fit a model to correct for remaining non-uniformity in stock solution concentration. For all 120 amplicon mixture samples in Figs. 2 and 3, we fit a model in which the amplicon concentration of each strain *i* was *C_i_* times greater than expected. We then calculate the mixture fractions that would have rcn equal.

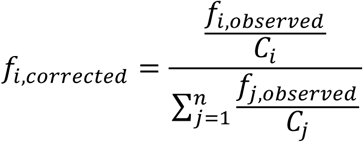

We estimated *C_i_* values by finding the *C_i_* that minimized the mean squared difference between the corrected fractions and the known fractions at which the amplicon stock solutions were volumetrically mixed. Arbitrarily, C_A_ was set to 1, and all C_i_ values were defined relative to that (therefore our model has seven free parameters). Concentration error estimates were as follows: C_B_=5.35 (not reliable due to poor estimates of f_observed_), C_C_=1.10, C_D_=0.80, C_E_=0.66, C_F_=0.91, C_G_=0.86, C_H_=0.75.

### Model community dynamics and Illumina sequencing

Communities contained non-overlapping sets of four marine isolates. Strains in each community were taxonomically classified using SINA^28^ and are listed below at the narrowest taxonomic level available. Communities are denoted by their subfigure label in Fig. 5.

A: G2R05 Cellulophaga, C3M06 Rhodobacteraceae, F3R02 Neptunomonas, G3M19 Celeribacter

B: I3R01 Vibrio, F3R08 Shewanella, E3M07 Paraglaciecola, A3R10 Tenacibaculum

C: A1M03 Alteromonas, G2M11 Colwellia, A1R15 Pseudoalteromonas, G2M05 Photobacterium

D: D3R06 Colwellia, E3M10 Cellulophaga, G2R14 Vibrio, I2M14 Marinobacterium

E: D2R05 Alteromonadaceae, E3R09 Winogradskyella, A1R03 Shewanella, D2R04 Rhodobacteraceae

F: G2M18 Saccharospirillaceae, I3M06 Marinobacterium, C2R09 Paracoccus, I2M19 Marinobacterium

G: C3R15 Flavobacteriaceae, E3R01 Tenacibaculum, B3M02 Psychromonas, G2R10 Vibrio

We grew communities at room temperature while shaking in 200uL of 2216 Marine Broth media in a 96-well deep well plate. Communities were diluted 100-fold and transferred to new plates every 24 hours, and sampled every 3 days for DNA extraction and sequencing. DNA extraction was performed using a Epicentre MasterPure Kit.

Illumina 16S V4-V5 library preparation and sequencing were performed at Integrated Microbiome Resource (IMR) on an Illumina MiSeq (paired-end, 300-basepair reads). For Sanger sequencing, samples were PCR-amplified using identical conditions as described for the amplicon mixtures (above), but in 25 μL volumes instead of 50 μL.

### Analysis of Illumina sequencing data

Next-generation sequencing of model communities resulted in 514,524 reads (average per sample of 64,316 reads, range over all samples of 46,633-72,095 reads). Paired-end reads were merged with vsearch –fastq_mergepairs (10 mismatches allowed in overlap region) and trimmed of primer sequences with cutadapt 1.16. We estimated read counts for each isolate by assigning trimmed and merged reads that perfectly matched a known isolate 16S rRNA V4-V5 sequence to that isolate. Reads that did not match a known isolate were discarded. Fractional abundances were estimated by dividing the read count for each isolate in a sample by the read counts for all isolates in that sample.

---

**Fig S1.**
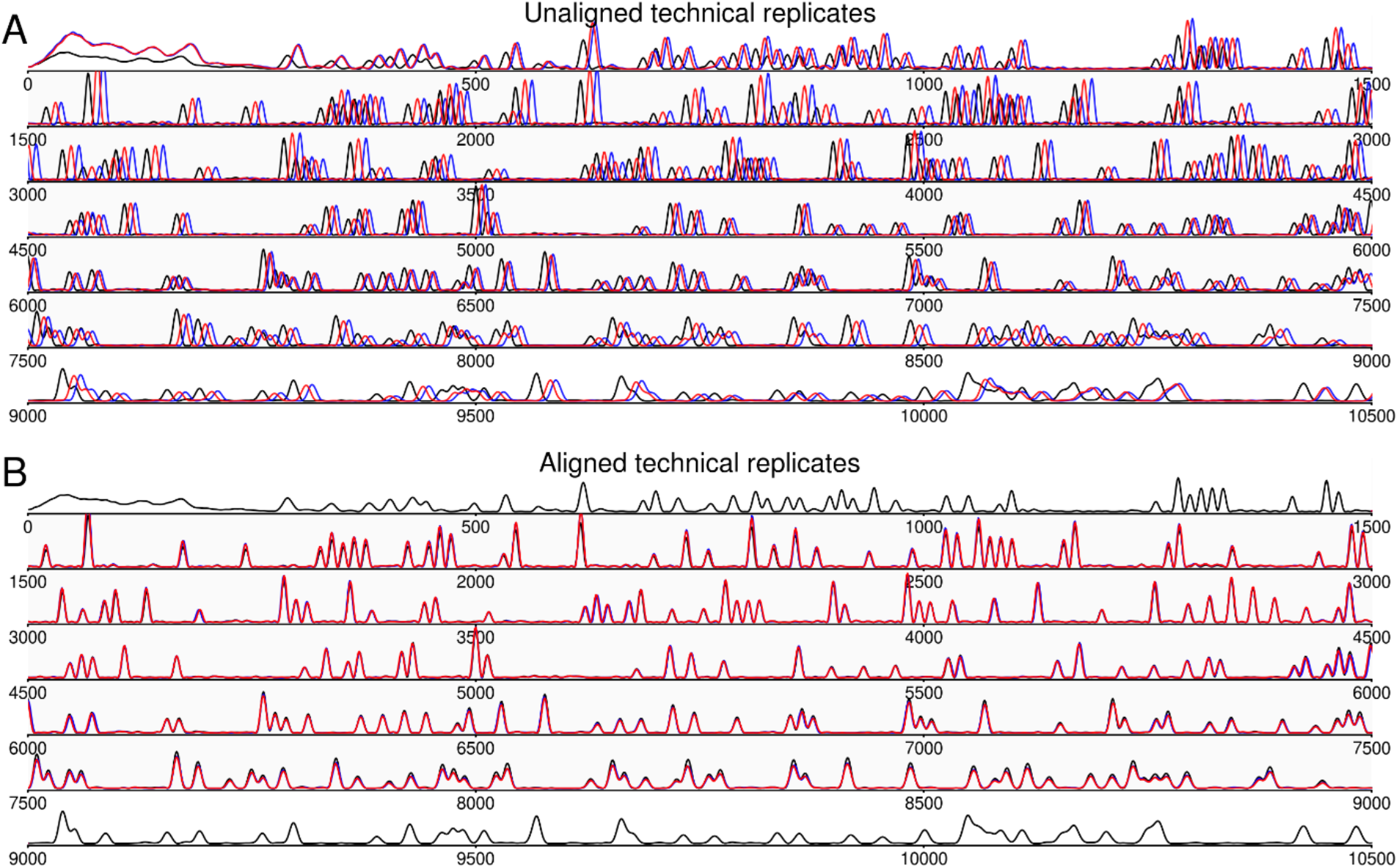
Aligning via time-warping can correct for temporal variability among technical replicates. (A) Three technical replicates (black, red and blue) for a sample of 16S DNA (showing only a single fluorescence channel for clarity). (B) Technical replicates two and three aligned to technical replicate one, over indices 1500-9000, covering ~630 bases.

**Fig S2.**
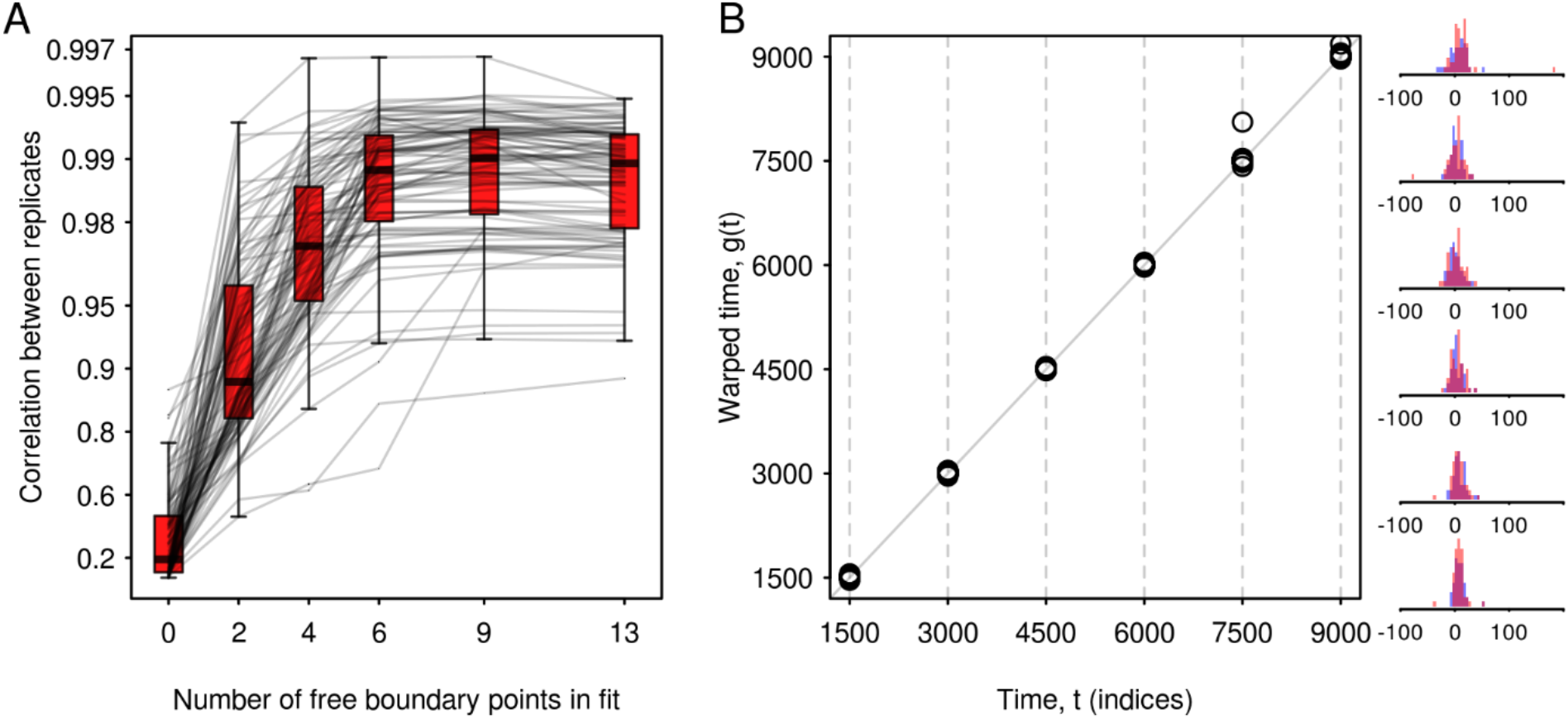
Determining optimal warping flexibility and typical range of warping parameters. (A) A six-parameter alignment yields good fits without unnecessary degrees of freedom. Fit quality was quantified as post-alignment Pearson correlation between technical replicates. With less than six boundary parameters, fits can be improved by increasing the warping flexibility, but beyond six parameters improvements are minimal. (B) Warping functions estimated from fitting each sample to its two technical replicates (N= 48 samples; 8 single-strain samples; 30 two-strain samples; 5 four-strain samples; 5 seven-strain samples). At right are histograms of offsets for each boundary (how many time indices the boundary was moved in the alignment). Pink bars show alignment parameters for replicate 2, blue bars show alignment parameters for replicate 3, suggesting no day-specific effects.

**Fig S3.**
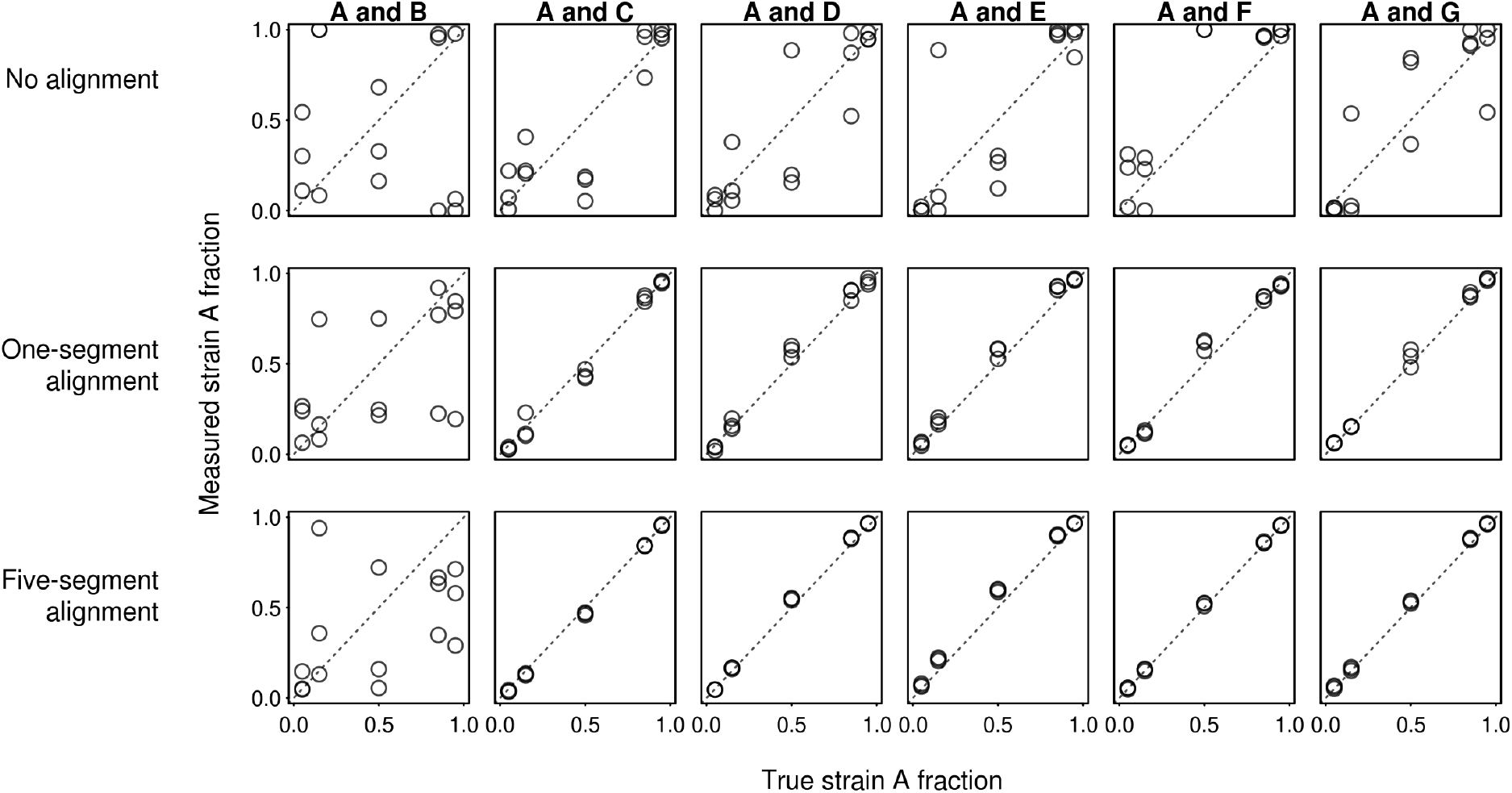
Flexible alignment is necessary to quantify community composition accurately. Without alignment, estimates of the fractions of each component are poor (open blue circles). With 2-parameter alignment it yields more accurate results (open red circles), but still much less precise than those obtained with 6-parameter alignment (black). Data in this figure is not corrected for error in stock solution concentrations.

**Fig S4.**
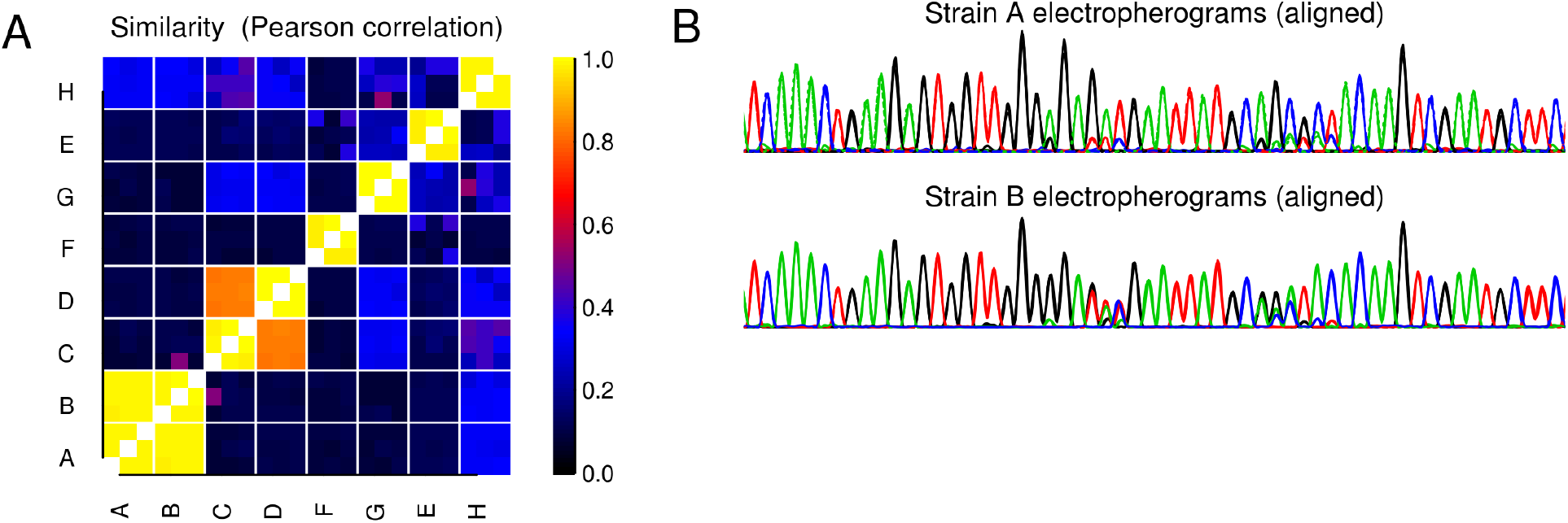
Strain similarity varies and is maximal between strains A and B. (A) Similarity matrix between single strains used in amplicon mixture analyses (Figs. 2-4), where the similarity metric is the Pearson correlation after alignment. Each amplicon was sequenced three times and therefore each 3×3 square shows correlations between all possible pairs of electropherograms. (B) After alignment, strains A and B are identical aside from two short stretches of ~5 bases each. Outside of the middle of the region shown (indices 4900-5600), the electropherograms are indistinguishable. Colors correspond to the four fluorescence channels.

**Fig S5.**
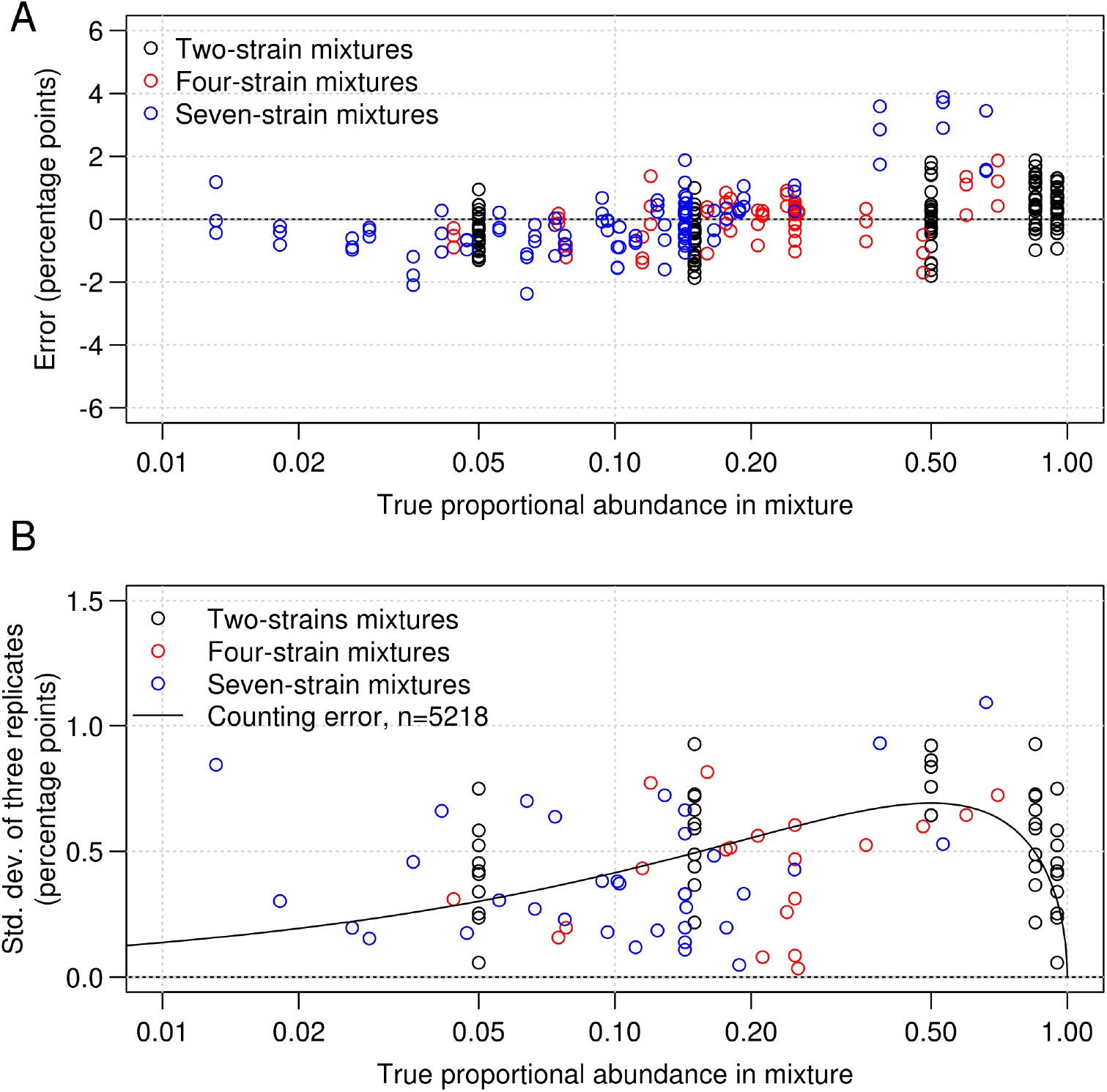
CASEU error magnitude is only weakly dependent on strain abundance. (A) Errors are similar in magnitude regardless of a strain’s abundance, though there is some bias in the seven-strain mixtures at higher proportions (blue circles). (B) Standard deviation of abundances calculated from triplicate Sanger sequencing measurements are generally around ~0.5% and are comparable to those that would be expected from counting based methods like NGS or plate counts with *n*=5218 counts (best fit to 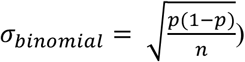.

**Fig S6.**
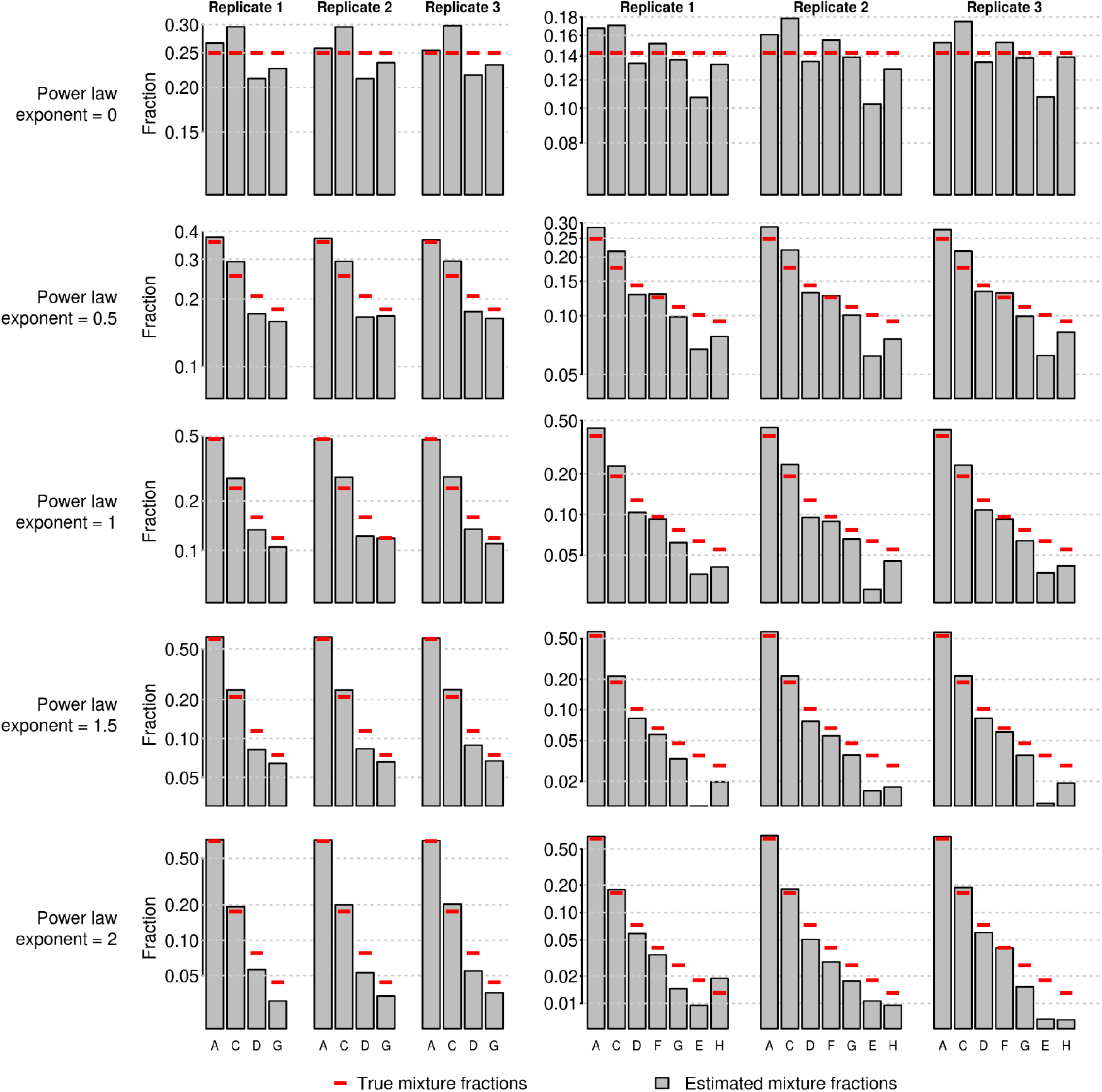
Without correcting for concentration errors in stock solutions, four- and seven-strain mixtures are consistent across replicates but biased. Data are the same as shown in Fig. 3 but without correcting for stock solution concentration errors.

**Fig S7.**
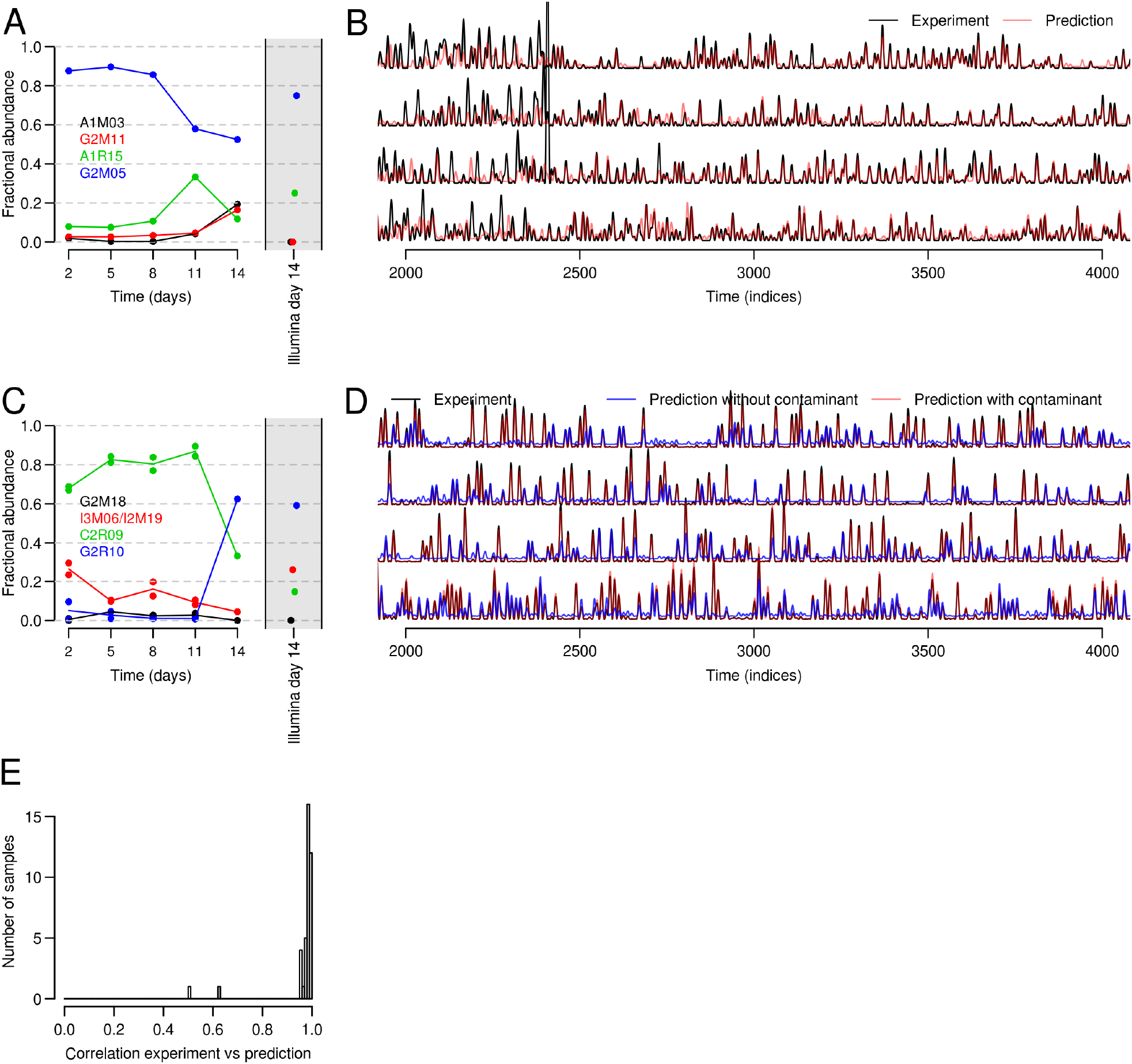
Assessing CASEU fits to detect sample preparation and/or sequencing errors. (A) Time course of a community for which the CASEU estimate for the last time point was not reliable and suggested problems with sequencing or sample preparation (below). (B) The CASEU fit to the day 14 sample (red lines) does not accurately reproduce the observed mixed electropherogram (black lines). Each of the four traces shows a single fluorescence channel of the electropherogram. The best fit remained poor even when excluding the spike around t=2400. (C) Time course of a community that was contaminated (contaminating strain in blue, G2R10) towards the end of the experiment. (Note also that this community was sequenced twice at the first four time points). (D) The CASEU fit to the day 14 community (blue) did not accurately reproduce the observed mixed electropherogram (black). After including the contaminating strain (identified by Illumina sequencing), the CASEU fit (red) reproduces the observed electropherogram. E) Histogram of correlations between experiment and prediction for the model communities (n=39), showing two clear outliers (the communities shown in (A) and (C)).

